# Chronic Pod-Mod E-Cigarette Aerosol Exposure Induces Aortic Dysfunction in Hypercholesterolemic Mice: Role of Oxidative Stress and Inflammation

**DOI:** 10.1101/2024.01.30.578110

**Authors:** Yasmeen M. Farra, Jacqueline Matz, Hannah Wilker, Hannah Kim, Cristobal Rivera, John Vlahos, Bhama Ramkhelawon, Jessica M. Oakes, Chiara Bellini

**Affiliations:** Department of Bioengineering, Northeastern University, Boston, USA; Division of Vascular and Endovascular Surgery, Department of Surgery, New York University Langone Medical Center, New York, USA; Department of Cell Biology, New York University Langone Medical Center, New York, USA

**Keywords:** Aortic Tissue Stiffness, Aortic Distensibility, Aortic Tissue Fibrosis, Endothelial Dysfunction, Inflammation, Oxidative Stress

## Abstract

**Objective:** Electronic (e-)cigarettes are the most used tobacco product amongst youth, and adult smokers favor e-cigarettes over approved cessations aids. Despite the lower perceived harm of vaping compared to smoking, inhalation of e-cigarette aerosol elicits cardiovascular responses that may lead to permanent injury when repeated over time. We thus aimed to infer the long-term outcomes of vaping on the function and structure of the aorta and shed light on the underlying cellular and molecular mechanisms.

**Approach and Results:** We exposed female hypercholesterolemic mice to either pod-mod e-cigarette aerosol or room air daily for 24 weeks. Chronic inhalation of e-cigarette aerosol triggered accumulation of inflammatory signals systemically and within aortic tissues, as well as T lymphocyte accrual in the aortic wall. Reduced eNOS expression and enhanced ROS production following eNOS uncoupling and NADPH oxidase activation curbed nitric oxide availability in the aorta of mice exposed to e-cigarette aerosol, impairing the endothelium-dependent vasodilatation that regulates blood flow distribution. Inhalation of e-cigarette aerosol thickened and stiffened aortic tissues via collagen deposition and remodeling, hindering the storage of elastic energy and limiting the cyclic distensibility that enables the aorta to function as a pressure reservoir. These effects combined contributed to raising systolic and pulse pressure above control levels.

**Conclusions:** Chronic inhalation of aerosol from pod-mod e-cigarettes promotes oxidative stress, inflammation, and fibrosis within aortic tissues, significantly impairing passive and vasoactive aortic functions. This evidence provides new insights on the biological processes that increase the risk for adverse cardiovascular events as a result of pod-mod e-cigarette vaping.

## 1. INTRODUCTION

Notwithstanding effective public health campaigns that have drastically curbed the prevalence of combustible cigarette smoking in the United States, 19% of adults and 13.4% of high school students have reported current use of commercial tobacco or nicotine products.^1, 2^ Classified as electronic nicotine delivery systems (ENDS), electronic cigarettes (or e-cigarettes) are battery-operated devices that serve to aerosolize a solution of propylene glycol and glycerin as a carrier for nicotine and flavorings. By incorporating nicotine salts as opposed to the harsh free-base form of the compound, the latest disposable and prefilled/refillable pod-mod devices generate aerosols that are easier to inhale^3, 4, 5^ and yet contain higher levels of nicotine compared to previous generation e-cigarettes.^6^ The popularity of these products has nearly doubled the likelihood of nicotine dependence in adolescent e-cigarette users,^7^ which in turn contributes to further reinforce emerging vaping habits.^1, 8, 9, 2^

Although vaping is often perceived as a safer alternative to cigarette smoking,^10, 11^ the physical properties and chemical composition of e-cigarette aerosol suggest otherwise. We have recently shown that e-cigarette devices emit levels of fine and ultra-fine particulate matter comparable to those found in cigarette smoke.^12^ Also similar to cigarette smoke, e-cigarette aerosol contains irritant aldehydes such as acrolein, free radicals, metals, and other chemicals that are harmful to human health.^5, 13, 14, 15^ While nervous system signaling due to airway irritation has been implicated in the immediate vascular response to inhaled smoke or aerosol,^16^ deposition of respirable particles into the deep lungs otherwise mediates the long-term outcomes.^17, 18^ Habitual e-cigarette users experience systemic oxidative stress,^19^ radial artery stiffening,^20^ and reduced brachial artery flow-mediated dilatation.^21^ Complementary to this evidence, endothelial nitric oxide synthase (eNOS) phosporylation^22^ and nitric oxide (NO) production^20^ are blunted in venous endothelial cells from e-cigarette users. Human umbilical vein endothelial cells (HUVECs) cultured with serum from e-cigarette users^21^ and human aortic endothelial cells (HAECs) incubated with e-liquids^22^ likewise secrete less NO. Exposure to e-cigarette aerosol extract^23^ and e-cigarette user serum^21^ further induces production of reactive oxygen species (ROS) in cultured HUVECs.

Corroborating epidemiological and *in vitro* evidence, chronic inhalation of aerosol from tank-style devices with free-base nicotine stiffens the common carotid artery^24^ and impairs the endothelium-dependent dilatation of the thoracic aorta^25, 24^ in wildtype mice. Reduced flow-mediated dilation of the femoral artery in live rats soon after exposure extends these findings to the peripheral vasculature and pod-mod e-cigarettes with nicotine salts.^26^ However, while activation of NADPH oxidases and eNOS uncoupling have been shown to enhance the oxidative burden in the aorta following inhalation of earlier generation e-cigarette aerosol,^27, 28^ the molecular mechanisms responsible for the adverse vascular outcomes of pod-mod vaping are poorly understood.^29^ Furthermore, the effects of prolonged exposure to pod-mod e-cigarette aerosol on microstructural composition, passive tissue mechanics, and vasoreactivity of elastic arteries demand additional consideration,^30^ to dissect the contribution of peripheral *vs.* central arterial stiffening on the cardio-vascular risk of vaping. Finally, the compounded role of hypercholesterolemia, which offers a substrate for oxidative stress, on the vascular outcomes of pod-mod vaping warrants further investigation. To this end, we characterized the effects of chronic pod-mod e-cigarette aerosol inhalation on the structure and function of the female Apoe*^−/−^* mouse aorta and explored the underlying cellular and molecular mechanisms. Our findings implicate chronic pod-mod vaping as a cardiovascular risk factor and provide evidence in support of regulatory guidance seeking to limit the hazards of chronic e-cigarette use over the coming decades.

## 2. METHODS

### Mouse exposure and weekly blood pressure

All experimental procedures involving live mice were approved by Northeastern University Institutional Animal Care and Use Committee (IACUC) and followed National Institutes of Health (NIH) guidelines. Adult female Apoe*^−/−^* mice were acquired from Jackson Laboratory (Bar Harbor, ME, USA) at 8±1 weeks of age and randomly assigned to either the e-cigarette aerosol or the air control exposure groups. Details on acclimation and exposure protocols have been reported elsewhere.^31, 12, 32^ Briefly, all mice were exposed for 95 minutes/day, 5 days/week, and over 24 weeks via nose-only aerosol delivery (SCIREQ; Montreal, QC, CA). The e-cigarette exposure group received aerosols from Virginia tobacco flavored JUULpods (JUUL Labs; San Francisco, CA, USA) containing 3% (35 mg/mL) nicotine and delivered at a rate of 2 puffs/min and puff volume of 55 mL/puff. The air control group inhaled filtered lab air. Peripheral blood pressure was measured weekly in awake and restrained mice (n=8 per group) using a non-invasive tail-cuff system (CODA, Kent Scientific; Torrington, CT, USA).

### Systemic inflammation and oxidative stress

Whole blood samples (n=20 per group) were obtained via submandibular collection at baseline (exposure week 0), week 8, week 16, and sacrifice (week 24). Following centrifugation, plasma was aspirated and stored at -80*^◦^*C. Samples were thawed on ice to analyze circulating levels of interleukins -2 (IL-2) and -6 (IL-6) with commercial ELISA assays (Quansys Biosciences; Logan, UT, USA). Additional blood samples collected at the 24-week endpoint (n=10 per group) were used to quantify the ratio of reduced (GSH) to oxidized (GSSG) circulating glutathione with a commercial ELISA kit (Abcam; Cambridge, UK) as an indicator of oxidative stress levels and redox imbalance.^33^

### Vasoactive biaxial testing

Deeply anesthetized (2-3% isoflurane) mice were euthanized by exsanguination the day after the final exposure. Whole aortas were immediately excised, immersed in chilled (4*^◦^*C) Hank’s buffered salt solution (HBSS), and prepared for mechanical testing as previously described.^31, 34^ Briefly, the ascending thoracic aorta (ATA) was sectioned between the aortic root and the brachiocephalic artery, while the suprarenal abdominal aorta (SAA) was segmented from the diaphragm to the right renal artery. Aortic samples were cleaned of perivascular tissue to facilitate ligation of lateral branches using a single strand of a braided 9-0 suture. Following cannulation onto glass micropipettes, aortic samples were transferred to a custom, computer-controlled biaxial mechanical device and immersed in warm (37*^◦^*C) Krebs-Ringer’s buffer solution. A 95% O_2_, 5% CO_2_ gas mixture was bubbled into the bath to maintain a pH of 7.4.

In preparation for active testing, SAA segments (n=8 per group) were acclimated for 10 minutes at luminal pressure of 80 mmHg and preferred axial stretch. Phenylephrine (L-Phenylephrine hydrochloride, 99%, Thermo Fisher Scientific; Waltham, MA, USA) was added to the bath at a concentration of 2*·*10*^−^*^6^ M to induce sub-maximal vasoconstriction over 15 minutes. Increasing doses of acetylcholine (Acetylcholine chloride, 99%, Thermo Fisher Scientific; Waltham, MA, USA) ranging from 10*^−^*^9^ to 10*^−^*^4^ M were administered sequentially through the pressurized luminal loop to probe for endothelium-dependent vasodilatation. At the end of the test, aortic samples were restored to their traction-free configuration and washed with fresh Krebs-Ringer’s buffer solution for 10 minutes. To assess the contribution of endothelial nitric oxide synthase (eNOS) signaling to the endothelium-dependent vasodilatation, L-NAME (NG-Nitroarginine methyl ester hydrochloride, Tocris Bioscience; Bristol, UK) at a 10*^−^*^4^ M concentration was administered luminally to pressurized and axially extended aortic samples and allowed to inhibit NOS activity over 30 minutes. Phenylephrine vasoconstriction followed by vasodilatation at increasing acetylcholine doses were then performed as described above. Aortic samples were once again equilibrated to their traction-free configuration and washed with fresh Krebs-Ringer’s buffer solution. Endothelium-independent vasodilatation was finally probed by administering increasing concentrations between 10*^−^*^9^ and 10*^−^*^4^ M of the nitric oxide (NO) donor sodium nitroprusside (MP Biomedicals; Irvine, CA, USA) through the lumen of pressurized, axially extended, and pre-constricted aortic samples. Sequential doses of either acetylcholine or sodium nitroprusside were administered 5 minutes apart to allow for the active response to stabilize. The outer diameter was imaged using a camera positioned in front of the bath, and the axial force was measured with a load cell coupled to the distal cannula, with both signals visualized within a custom LabView (National Instrument; Austin, TX, USA) interface throughout active testing. Experimental data was analyzed to extract the dose response of each aortic sample to either acetylcholine or sodium nitroprusside. Percent vasoconstriction to phenylephrine was calculated as (*od_base_ − od_phen_*)*/od_base_ ·* 100, where *od_base_* is the steady-state outer diameter prior to addition of any chemical, and *od_phen_* is the outer diameter 15 minutes after phenylephrine administration but prior to the addition of any acetylcholine or sodium nitroprusside. As diameter varied across aortic samples, percent vasodilatation was normalized to the maximal diameter change as (*od_dose_ − od_phen_*)*/*(*od_base_ − od_phen_*) *·* 100, where *od_dose_* is the steady-state outer diameter at each dose of either acetylcholine or sodium nitroprusside. Finally, the difference between the percent vasodilatation at the highest dose of acetylcholine alone and after L-NAME administration was calculated as an estimate of NO bioavailability.

### Passive mechanical testing

The biaxial testing device that facilitated vasoactive testing was also used to characterize the passive mechanical response of aortic samples (n=10 e-cigarette and n=16 air control). Details on the passive inflation/extension testing protocols and analysis are provided elsewhere.^31^ Briefly, ATA and SAA segments extended to their preferred axial stretch were first acclimated then preconditioned by cyclic luminal pressurization while submerged in room temperature HBSS. Following re-estimation of the traction free configuration and preferred axial stretch, aortic samples were subjected to three pressure– diameter tests (cyclic luminal pressurization between 10 and 140 mmHg while extended at or *±*5% the preferred axial stretch) and four axial force–length tests (cyclic axial stretching to reach a set axial force while maintained under luminal pressure of either 10, 60, 100, or 140 mmHg). A nonlinear regression algorithm was implemented to fit the experimental data to a microstructurally-motivated four-fiber family strain energy potential (*W*) in the form

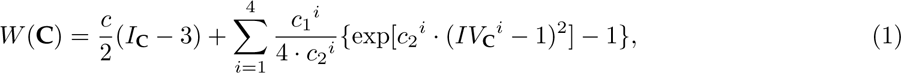

where the first isotropic Neo-Hookean term (coefficient *c* with the dimension of a stress) accounts for the contribution of elastic fibers and amorphous matrix, while the second term (coefficients *c*_1_*^i^* with the dimension of a stress and unitless coefficients *c*_2_*^i^*) models collagen and smooth muscle bundles as anisotropic Fung-type solids^35^ along the axial (*i* = 1), circumferential (*i* = 2), and two symmetric diagonal (*i* = 3, 4) directions. *I***_C_** and *IV***_C_***^i^* are the first and fourth invariants of the right Cauchy-Green deformation tensor **C**, respectively. Best-fit material parameters for Equation 1 were used to predict geometry and intrinsic tissue properties at group-specific systolic pressure and sample-specific preferred axial stretch. The circumferential and axial components of linearized tissue stiffness were calculated using the small-on-large approach.^36^ Finally, the structural stiffness of the aorta as a conduit was expressed in terms of distensibility (*id_sys_ −id_dias_*)*/*(*P_sys_ −P_dias_*), where *id* is the luminal diameter, *P* is the luminal pressure, and the subscripts *sys* and *dias* refer to systole and diastole, respectively.

### Histology and immunohistochemistry

Following mechanical testing, a subset of aortic samples (n=6 ATA and n=6 SAA per group) were fixed overnight in 10% neutral-buffered formalin and stored in 70% ethanol at 4*^◦^*C. Fixed tissues were embedded in paraffin wax and cut into serial cross-sections with thickness set to 5*µ*m and starting at *∼*500*µ*m beyond the proximal end of each aortic segment. A first cohort of tissue sections were stained with MOVAT’s pentachrome and picrosirius red histological stains and imaged at 40x magnification under brightfield and polarized light, respectively (DM4 B upright microscope, Leica Microsystems; Wetzlar, Germany). The area occupied by microstructural constituents was automatically quantified by thresholding digitalized images for values of pixel hue, saturation, and lightness according to previously reported protocols.^31^ Chromogenic immunostaining for COL1A1 (1:500; ab34710, Abcam; Cambridge, UK), COL3A1 (1:1500; 22734-1-AP, Proteintech; Wuhan, China), CD68 (1:500; 97778, Cell Signaling Technology; Danvers, MA, USA), and VCAM-1 (1:500; ab134047, Abcam; Cambridge, UK) was executed on a second cohort of tissue sections. Imaging and analysis were performed as described above to quantify the area fraction occupied by positively-stained tissues.

A separate set of ATA samples (n=4 per group) were immersed in optimal cutting temperature compound (OCT; 4585, Thermo Fisher Scientific; Waltham, MA, USA) immediately after excision and frozen at 80*^◦^*C for fluorescent immunostaining. Frozen tissues were sectioned into 7-*µ*m-thick slices and incubated overnight with anti-eNOS (1:100; ab76198, Abcam; Cambridge, UK), anti-PECAM1 (1:200; 14-0311-82, Invitrogen; Carlsbad, CA, USA), anti-ACTA2 (1:400; 48938S, Cell Signaling; Danvers, MA, USA), anti-IL6 (1:100; ab179570, Abcam; Cambridge, UK), and anti-CD4 (1:200; 14-0042-82, Invitrogen; Carlsbad, CA, USA) primary antibodies. Alexa Fluor 488 and 568 conjugated goat anti-rabbit or anti-rat IgG antibodies (1:500 each; Invitrogen; Carlsbad, CA, USA) were used for fluorescent signal detection. 4’,6-diamidino-2-phenylindole (DAPI, 1:5000; Invitrogen; Carlsbad, CA, USA) allowed visualization of cell nuclei. Images were acquired at 40x magnification (LSM 710 confocal microscope and ZEN software, Carl Zeiss; Oberkochen, Germany). Identical acquisition parameters were set to capture tissue sections from e-cigarette and air control mice. Image thresholding for CD4+ positive cells and mean fluorescent intensity per unit area for all other stains were computed using ImageJ software (NIH; Bethesda, MD, USA).

### Atherosclerotic lesion morphology

Atherosclerotic lesions were first isolated between the luminal edge and the internal elastic lamina of aortic root sections that had been either stained with MOVAT’s Pentachrome and picrosirius red stains or incubated with anti-CD68 antibodies for chromogenic immunostaining. Regions that were devoid of cells but featured cellular debris and cholesterol crystals were classified as necrotic cores.^37^ Collagen rims surrounding necrotic cores were considered as fibrous caps. Total lesion area, area occupied by collagen within the lesion, percent of the lesion area occupied by the necrotic core, average fibrous cap thickness, and percent of the lesion area positive for CD68 were measured following protocols reported elsewhere^31^ to estimate the atherosclerotic burden.

### Detection of reactive oxygen species in aortic tissues

Dihydroethidium hydroethidine (DHE) was used as a probe to detect reactive oxygen species (ROS) *in situ*. DHE is a blue fluorescent dye that interacts with superoxide anion to form oxyethidium, which in turns intercalates with nucleic acids and emits red fluorescence detectable by microscopy.^38^ Frozen tissue sections from ATA samples (n=4 per group) were stained with DHE (10*µ*M; D1168, Invitrogen; Carlsbad, CA, USA) in a dark humidified incubator at 37*^◦^*C for 20 minutes. To gauge the contribution of uncoupled eNOS to ROS production in the aorta, a subset of tissue sections were incubated for 30 minutes at 37*^◦^*C with the NOS inhibitor L-NAME (500 *µ*M; Tocris Bioscience; Bristol, UK) prior to DHE staining. To further evaluate the role of NADPH oxidase activation, a separate cohort of tissue sections were pre-incubated for 30 minutes at 37*^◦^*C with either the NOX1 inhibitor ML171 (1 *µ*M; CAS 6631-94-3, Sigma Aldrich; St. Louis, MO, USA) or the NOX2 inhibitor GSK (50 *µ*M; GSK2795039, Sigma Aldrich; St. Louis, MO, USA). Tissue sections were imaged immediately following staining at 40x magnification to detect red DHE fluorescence (LSM 710 confocal microscope and ZEN software, Carl Zeiss; Oberkochen, Germany).

### Gene expression via RT-qPCR

Whole mouse aortas (n=18 per group) cleaned of perivascular tissue were perfused with sterile HBSS and stabilized overnight at 4*^◦^*C in RNAProtect reagent (Qiagen; Hilden, Germany) before long-term storage at -80*^◦^*C. Frozen aortic samples were finely ground into powder using a pre-chilled mortar and pestle to purify RNA using the RNeasy mini kit as per manufacturer instructions (Qiagen; Hilden, Germany). Quantity and quality of RNA was measured using a Nanodrop ND-1000 spec-trophotometer (Thermo Scientific; Waltham, MA, USA). RNA concentrations were normalized and reverse transcription was performed using the RNA to cDNA EcoDry premix kit (Takara Bio USA; San Jose, CA, USA). Taqman universal master mix and Taqman probes (Thermo Fisher Scientific; Waltham, MA, USA) were used for amplification and detection of specific expression of *Col1a1*, *Col3a1*, *Eln*, *Tgfb1*, *Timp1*, *Nos3*, and *Il6*, with *Actb* as the housekeeping gene. Quantitative real-time PCR reactions were performed in triplicate using a QuantStudio 5 Real-Time PCR system (Applied Biosystems; Foster City, CA, USA). Cycle threshold (C*_t_*) values were analyzed and fold-change gene expression over the housekeeping gene was calculated by the comparative cycle method (2*^−^*^ΔΔ*Ct*^).

### Statistics

Unless otherwise specified, data are presented as means *±* SE. Unpaired Student’s t tests with a two-tailed distribution and heteroscedastic variance assumption facilitated the comparison of single-metric data between the e-cigarette and air control groups. Two-way ANOVA was used to assess the effect of treatment (e-cigarette aerosol or air control inhalation) and duration of exposure (weeks) for longitudinal measurements, with Bonferroni correction for multiple comparisons. Differences were considered statistically significant if p *<* 0.05.

## 3. RESULTS

### E-cigarette aerosol exposure raised systolic and pulse pressure

Weekly noninvasive measurements of peripheral blood pressure allowed for longitudinal monitoring of cardiovascular function in e-cigarette and air control mice. Systolic pressure progressively increased from week 1 to 16 then stabilized to endpoint in mice exposed to pod-mod e-cigarette aerosol, while it fluctuated around baseline in air controls (Figure 1A). Following 24 weeks of exposure, systolic blood pressure was significantly higher in e-cigarette mice (127 *±* 1 mmHg) compared to air controls (110 *±* 2 mmHg). Diastolic pressure remained consistent throughout exposure in both groups (data not shown), and no difference emerged between mice exposed to e-cigarette aerosol (86 *±* 2 mmHg) and air controls (84 *±* 1 mmHg) at endpoint. As a result, the average pulse pressure increased from 26 *±* 1 mmHg at baseline to 37 *±* 1 mmHg by the end of exposure in the e-cigarette group, while it did not change significantly in air controls (Figure 1B).

**Figure 1.**
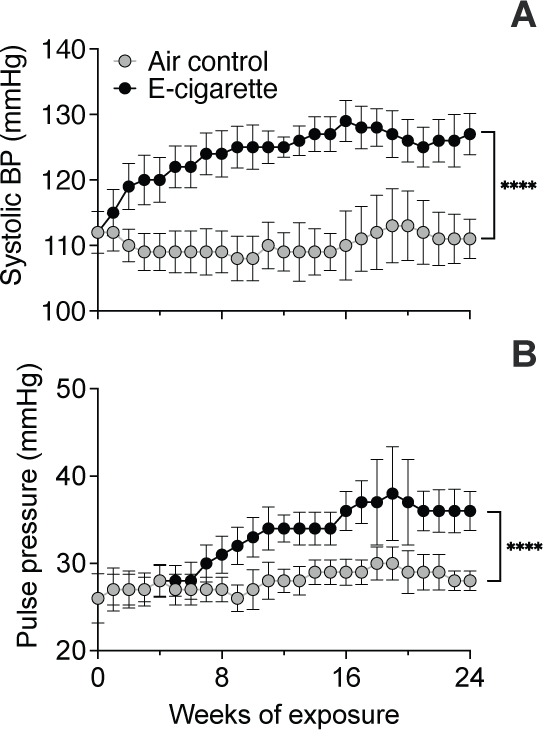
Weekly blood pressure measurements in air control and e-cigarette mice throughout the study. Average systolic (A) and pulse (B) pressure in mice exposed to e-cigarette aerosol compared to air control. Chronic inhalation of pod-mod e-cigarette aerosol progressively increased systolic and pulse pressure. Statistical significance denoted by **** for p *<* 0.0001 in e-cigarette *vs*. air control.

### E-cigarette aerosol exposure impaired endothelium-dependent and nitric oxide (NO)-mediated aortic dilatation

Evidence of impaired flow-mediated dilatation in e-cigarette users compared to nonusers^21^ motivated evaluating the vasoactive and endothelial function of the aorta in mice exposed to pod-mod e-cigarette aerosol for 24 weeks. The submaximal vasoconstriction of the aorta upon adrenergic stimulation with phenylephrine was comparable between e-cigarette and air control mice (e-cigarette: *−*23 *±* 1 %, air control: *−*21 *±* 1 % relative change in diameter). Similarly, exposure to e-cigarette aerosol did not affect the relative dilatation of the aorta in response to sodium nitroprusside with respect to air control (Figure 2A, Table S1). Nonetheless, chronic inhalation of e-cigarette aerosol reduced the relative change in aortic diameter following administration of acetylcholine, compared to air control (circles, Figure 2B, Table S1). Furthermore, pre-treatment with nitric oxide synthase (NOS) inhibitor L-NAME attenuated the response to acetylcholine in the air control but not the e-cigarette aorta (diamonds, Figure 2B, Table S1). Abrogation of the endothelium-dependent and NO-mediated dilatation allowed for indirect assessment of reduced NO availability in the e-cigarette aorta against air control (e-cigarette: -4 *±* 6%, air control: 34 *±* 11%; Figure 2C). We therefore considered potential sources of NO depletion and noted downregulation of protein (Figure 2D-E) and mRNA expression (Figure 2F) for endothelial nitric oxide synthase (eNOS) in the aorta of e-cigarette mice contrasted to air control.

**Figure 2.**
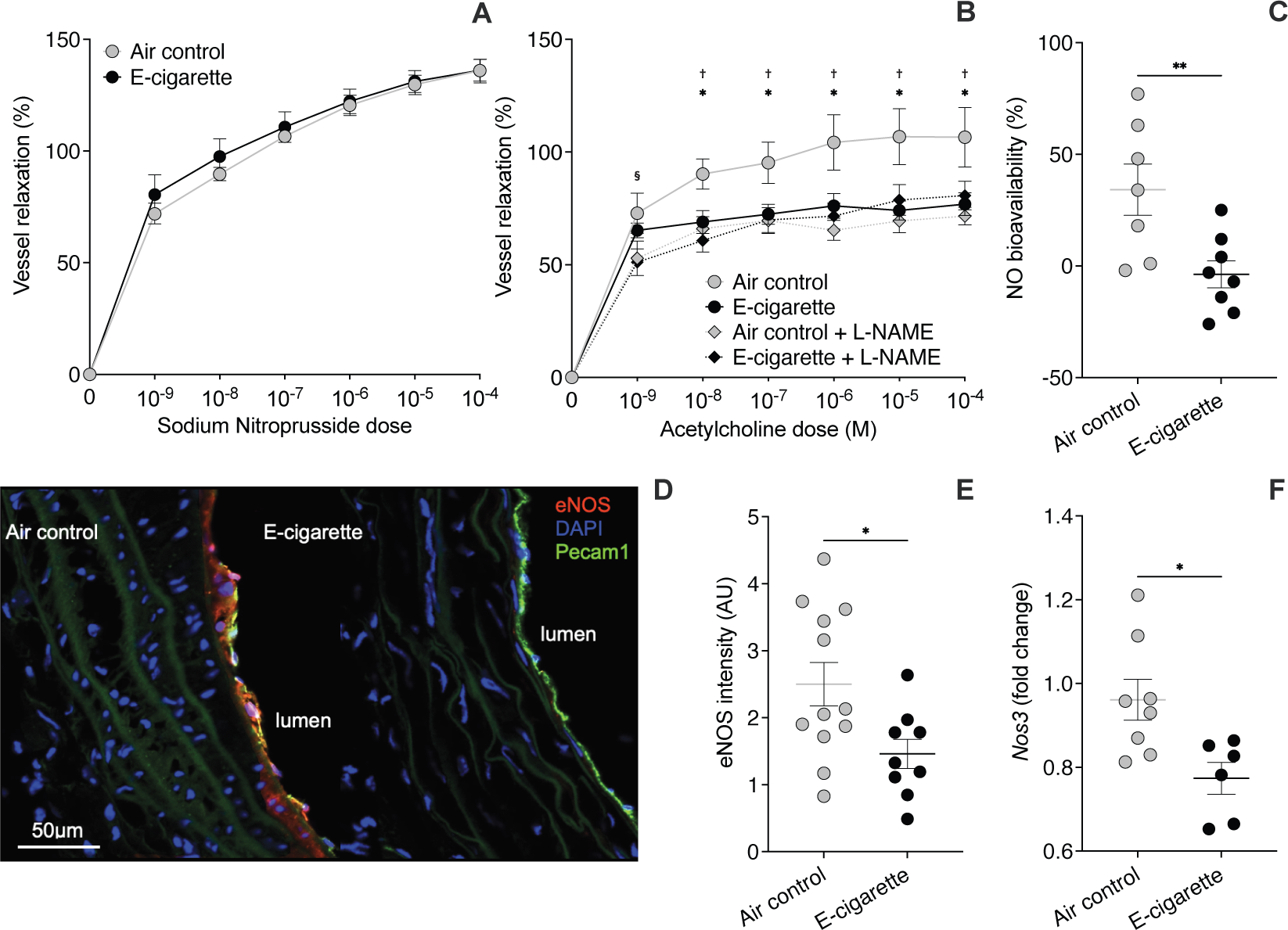
Vasodilator response of the aortic wall and eNOS expression in aortic tissues from air control and e-cigarette mice. The percent recovery of outer diameter from vasoconstriction by submaximal phenylephrine dose and in response to increasing concentrations of sodium nitroprusside (NO donor) measures the endothelium-independent vasodilatation (A). The percent recovery of outer diameter from vasoconstriction by submaximal phenylephrine dose and in response to increasing concentrations of acetylcholine measures the endothelium-dependent vasodilatation (B, circles). Incubation of aortic samples with L-NAME (NOS inhibitor) ahead of phenylephrine vasoconstriction and acetylcholine treatment quantifies the portion of endothelium-dependent vasodilatation that does not rely upon NO release (B, diamond). The difference between the dilatation to the largest acetylcholine dose without and with L-NAME provides an indirect estimate of NO availability in aortic tissues (C). Expression of eNOS in endothelial cells (E) from immunostaining (red) and colocalization with Pecam1 (green) in aortic cross-sections (D). mRNA expression of the *Nos3* gene coding for eNOS in whole aorta tissue samples (F). Chronic exposure to pod-mod e-cigarette aerosol hindered the endothelium-dependent vasodilatation of the aortic wall via abrogation of the NO-mediated response. Statistical significance denoted by * for p*<*0.05 or ** for p*<*0.01 in näıve e-cigarette *vs*. air control, † for p*<*0.05 in L-NAME pre-treated *vs*. näıve in air control, and § for p*<*0.05 in L-NAME pre-treated *vs*. näıve in e-cigarette.

### E-cigarette aerosol exposure elicited production of reactive oxygen species (ROS) in the aortic wall and systemic oxidative stress

Impairment of the NO-mediated contribution to the endothelium-dependent vasodilatation of the e-cigarette aorta compelled assessing the production of reactive oxygen species that may react with NO to form peroxynitrates. Fluorescent signal from dihydroethidium (DHE) oxidation products in excess of air control levels implicated inhalation of pod-mod e-cigarette aerosol in the production of superoxide anion within the the aortic wall (Figure 3A-B,F). Pre-incubation with NOS inhibitor L-NAME and NOX2 inhibitor GSK reduced fluorescence in the e-cigarette aorta to air control values, while pre-treatment with NOX1 inhibitor ML171 did not affect DHE oxidation (Figure 3C-E,F). Alongside local ROS accumulation in the aorta, exposure to e-cigarette aerosol lowered the GSH:GSSG ratio in blood plasma, aggravating systemic oxidative burden with respect to air control conditions (Figure 3G).

**Figure 3.**
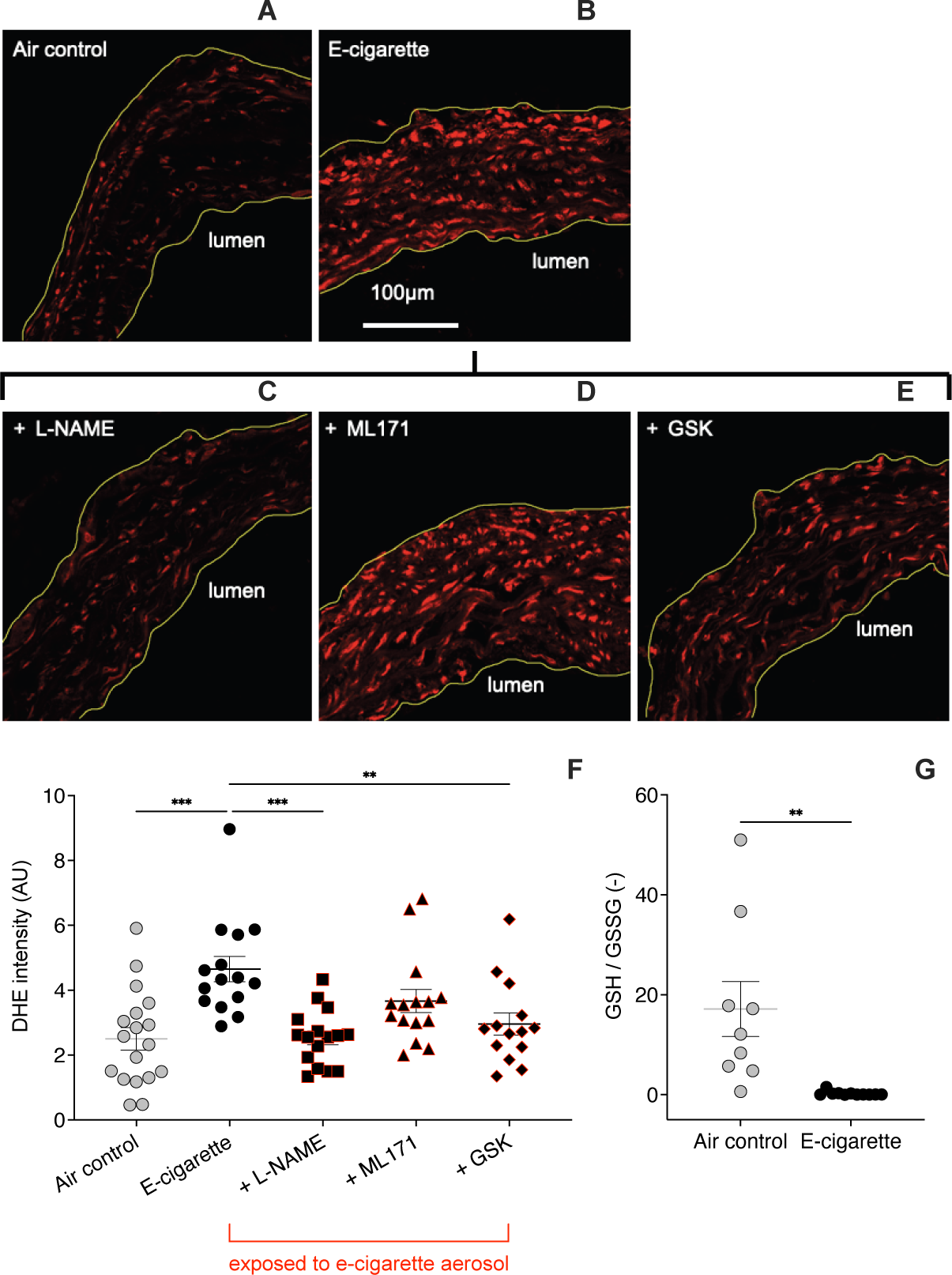
Systemic and local oxidative burden in aortic tissues from air control and e-cigarette mice. The release of (red) fluorescent products upon oxidation of dihydroethidium (DHE) staining by superoxide anion facilitates estimation of oxidative stress levels (F) within aortic tissues of e-cigarette (B) and air control (A) mice. Pre-incubation of aortic cross-sections with either the NOS inhibitor L-NAME (C), the NOX1 inhibitor ML171 (D), or the NOX2 inhibitor GSK (E) ahead of DHE staining weighs the contribution of established superoxide anion sources to the oxidative burden in aortic tissues (F). The ratio of reduced (GSH) to oxdized (GSSG) glutathione in the mouse blood plasma offers a marker of systemic oxidative stress (G). Superoxide production by uncoupled e-NOS and NOX2 fueled oxidative stress in aortic tissues of mice exposed to pod-mod e-cigarette aerosol. Statistical significance denoted by ** for p *<* 0.01 or *** for p *<* 0.001 in e-cigarette *vs*. air control.

### E-cigarette aerosol exposure stiffened the aorta and hindered the Windkessel function

Sustained increase in the structural stiffness of central arteries has been reported for up to 1 hour following e-cigarette vaping in otherwise healthy smokers^39^ and occasional smokers.^40^ Given the prognostic value of aortic stiffening, we evaluated this metric in näıve mice exposed to pod-mod e-cigarette aerosol, and further dissected contributing geometrical and tissue mechanical parameters. We show that chronic inhalation of e-cigarette aerosol altered the passive biaxial mechanical response of tissues from the ascending thoracic (ATA) and suprarenal abdominal (SAA) aorta as described by best-fit material parameters in Table S2.

The average circumferential stress *vs.* stretch curve of aortic tissues from mice exposed to e-cigarette aerosol extended to the right of air controls (Figure 4 A-B) to accommodate the larger systolic pressure (Figure 1A) while preserving the circumferential stress in both ATA (e-cigarette: 221 *±* 20 kPa, air control: 250 *±* 14 kPa; Figure 4E, Table S3) and SAA (e-cigarette: 223 *±* 7 kPa, air control: 223 *±* 9 kPa; Figure 4E, Table S3) segments. However, SAA tissues from e-cigarette mice exhibited higher stiffness (e-cigarette: 2.76 *±* 0.23 MPa, air control: 1.95 *±* 0.15 MPa; Figure 4F, Table S3) and experienced larger circumferential stretch (e-cigarette: 1.68 *±* 0.03, air control: 1.55 *±* 0.02; Figure 4G, Table S3) than controls at group-specific systolic pressure. Tissues from the ATA followed the same trend in circumferential stiffness (e-cigarette: 2.01 *±* 0.22 MPa, air control: 1.73 *±* 0.12 MPa; Figure 4F, Table S3) and stretch (e-cigarette: 1.80 *±* 0.12, air control: 1.68 *±* 0.04; Figure 4G, Table S3), though the differences between the two exposure groups did not attain statistical significance. Paralleling the larger pulse pressure (Figure 1B) with e-cigarette aerosol against air exposure, the cyclic circumferential stretch was higher in both ATA (e-cigarette: 1.087 *±* 0.003, air control: 1.066 *±* 0.003) and SAA (e-cigarette: 1.056 *±* 0.003, air control: 1.046 *±* 0.002) tissues, suggesting that the e-cigarette aorta exhibited greater dynamic range throughout the cardiac cycle (Figure 4H).

**Figure 4.**
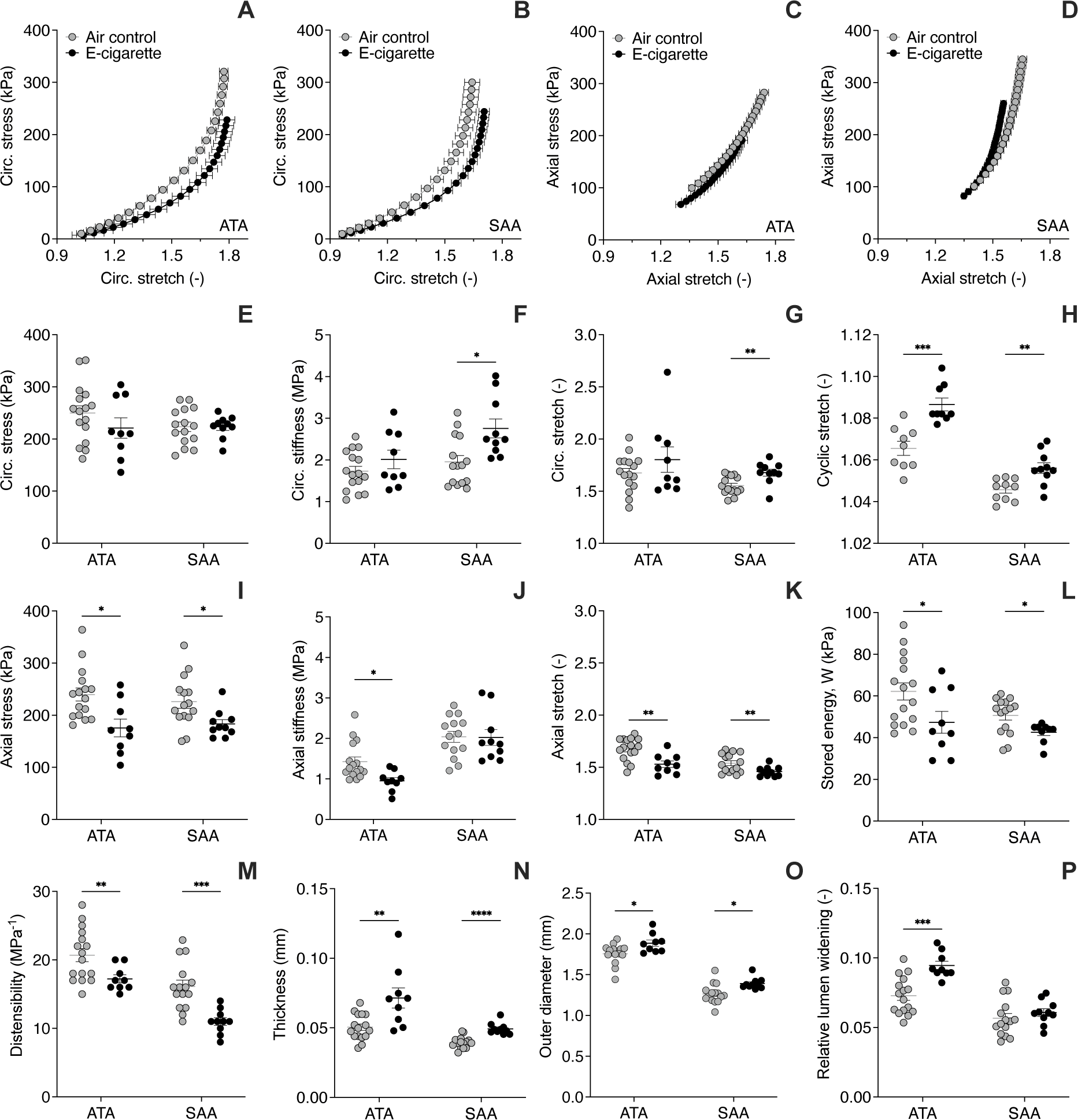
Geometrical, structural, and mechanical properties of the aortic wall and aortic tissues from air control and e-cigarette mice. Average global mechanical behavior of aortic tissues in the circumferential (A-B) and axial (C-D) directions in response to applied experimental loads. Estimated descriptors of biaxial aortic tissues properties (E-L), structural attributes of the aorta as a conduit (M), and aortic wall geometry (N-P) under *in vivo* loads. Mechanical outcomes are reported for the ascending thoracic (ATA) and suprarenal abdominal (SAA) aorta. Metrics of biaxial stress (E, I), biaxial stiffness (F, J), circumferential stretch (G), as well as energy (L), thickness (N), and outer diameter (O) are calculated at group-specific systolic pressure and predicted *in vivo* axial stretch (K). Cyclic stretch (H) measures the circumferential deformation of aortic tissues from diastole to systole. Distensibility (M) normalizes the relative widening of the aortic lumen (P) between diastole and systole to the group-specific pulse pressure. Thickening and stiffening of aortic tissues as a result of chronic pod-mod e-cigarette aerosol inhalation reduced their capacity for energy storage and limited the distension and recoil of the aortic wall throughout the cardiac cycle. Statistical significance within the same region denoted by * for p *<* 0.05, ** for p *<* 0.01, *** for p *<* 0.001, and **** for p *<* 0.0001 in e-cigarette *vs*. air control.

The axial stress *vs.* stretch response of aortic tissues shifted leftward and downward upon exposure to ecigarette aerosol, contrasted to air control (Figure 4 C-D). Lower systolic stiffness (e-cigarette: 0.95 *±* 0.08 MPa, air control: 1.43 *±* 0.11 MPa; Figure 4J, Table S3) and axial stretch (e-cigarette: 1.53 *±* 0.03, air control: 1.68 *±* 0.03; Figure 4K, Table S3) contributed to reduce the axial stress in ATA tissues from e-cigarette mice below air control levels (e-cigarette: 176 *±* 17 kPa, air control: 240 *±* 12 kPa in control; Figure 4I, Table S3). Axial stress (e-cigarette: 183 *±* 8 kPa, air control: 226 *±* 13 kPa; Figure 4I, Table S3) and stretch (e-cigarette: 1.46 *±* 0.02, air control: 1.54 *±* 0.02; Figure 4K, Table S3) in SAA tissues from mice exposed to e-cigarette aerosol were also lower than air controls, though no significant difference in axial stiffness was noted between the two groups (e-cigarette: 2.02 *±* 0.19 MPa, air control: 2.04 *±* 0.14 MPa; Figure 4J, Table S3). The axial stretch decline following e-cigarette aerosol inhalation further limited the systolic elastic energy stored within ATA (e-cigarette: 47 *±* 5 kPa; air control: 62 *±* 4 kPa) and SAA (e-cigarette: 43 *±* 2 kPa, air control: 51 *±* 2 kPa; Figure 4L, Table S3) tissues contrasted to air controls.

Chronic exposure to e-cigarette aerosol prompted thickening of the ATA (e-cigarette: 71 *±* 7 *µ*m, air control: 50 *±* 2) and SAA (e-cigarette: 49 *±* 1 *µ*m, air control: 40 *±* 1 *µ*m) wall above air control values (Figure 4N, Table S3). Compounded by higher systolic pressure (Figure 1A), the outward remodeling lead to larger adventitial diameters in both ATA (e-cigarette: 1885 *±* 39 *µ*m, air control: 1757 *±* 30 *µ*m) and SAA (e-cigarette: 1391 *±* 21 *µ*m, air control: 1265 *±* 31 *µ*m) segments (Figure 4O, Table S3). Consistent with greater cyclic stretch following e-cigarette exposure, the relative change in luminal diameter between diastole and systole was larger in the aorta of e-cigarette mice compared to air controls, although the difference reached statistical significance in the ATA (e-cigarette: 0.095 *±* 0.003, air control: 0.073 *±* 0.003) but not the SAA (e-cigarette: 0.061 *±* 0.003, air control: 0.057 *±* 0.003) segment (Figure 4P). Accounting for the rise in pulse pressure with e-cigarette aerosol inhalation, however, revealed compromised cyclic distensibility of ATA (e-cigarette: 17 *±* 1 MPa*^−^*^1^, air control: 21 *±* 1 MPa*^−^*^1^) and SAA (e-cigarette: 11 *±* 1 MPa*^−^*^1^, air control: 16 *±* 1 MPa*^−^*^1^) segments in the e-cigarette group against air controls (Figure 4M, Table S3).

### E-cigarette aerosol exposure promoted aortic wall thickening due to collagen deposition and remodeling

Stiffening and thickening of the aortic wall prompted analyzing the microstructural composition of aortic tissues from e-cigarette mice, to shed light on the remodeling process that supported noted functional shifts. In agreement with wall thickness measurements on fresh tissues (Table S3), the area of histological cross-sections obtained from the aorta of mice exposed to pod-mod e-cigarette aerosol was larger than air control in ATA (e-cigarette: 5.88 *±* 0.12 mm^2^, air control: 5.10 *±* 0.27 mm^2^) and SAA (e-cigarette: 2.88 *±* 0.13 mm^2^, air control: 2.48 *±* 0.04 mm^2^) tissues (Table S4).

The total area occupied by collagen in aortic cross sections stained with picrosirius red and imaged under polarized light (Figure 5A) was more extensive following e-cigarette aerosol exposure than air control in ATA (e-cigarette: 2.09 *±* 0.14 mm^2^, air control: 1.56 *±* 0.06 mm^2^) and SAA (e-cigarette: 1.30 *±* 0.05 mm^2^, air control: 1.04 *±* 0.03 mm^2^) tissues (Figure 5B, Table S4). Segmented by layer, the adventitial collagen area was broader in both regions of the e-cigarette aorta compared to air control, while the trend toward larger medial collagen area was significant in the ATA alone (Table S4). The percent of cross-sectional area that stained positive for COL1A1 (Figure 5C) increased following exposure to e-cigarette aerosol in ATA (ecigarette: 26.7 *±* 2.1%, air control: 16.2 *±* 0.6%) and SSA (e-cigarette: 45.2 *±* 0.8%, air control: 32.7 *±* 3.8%) tissues (Figure 5D). However, the COL3A1-positive area (Figure 5E) covered a larger percentage of the wall in the ATA (e-cigarette: 34.0 *±* 3.3%, air control: 17.3 *±* 1.6%) but not the SAA (e-cigarette: 20.1 *±* 1.2%, air control: 21.1 *±* 4.2%) region of the e-cigarette aorta, contrasted with air control (Figure 5F). Given regional difference in collagen remodeling, *Col1a1* but not *Col3a1* gene expression was upregulated in whole aorta tissue samples from mice exposed to e-cigarette aerosol compared to air control (Figure 5G). Upregulation of *Tgfb1* (Figure 5H) and *Timp1* (Figure 5I) gene expression further evidenced pro-fibrotic signaling in the e-cigarette aorta with respect to air control.

**Figure 5.**
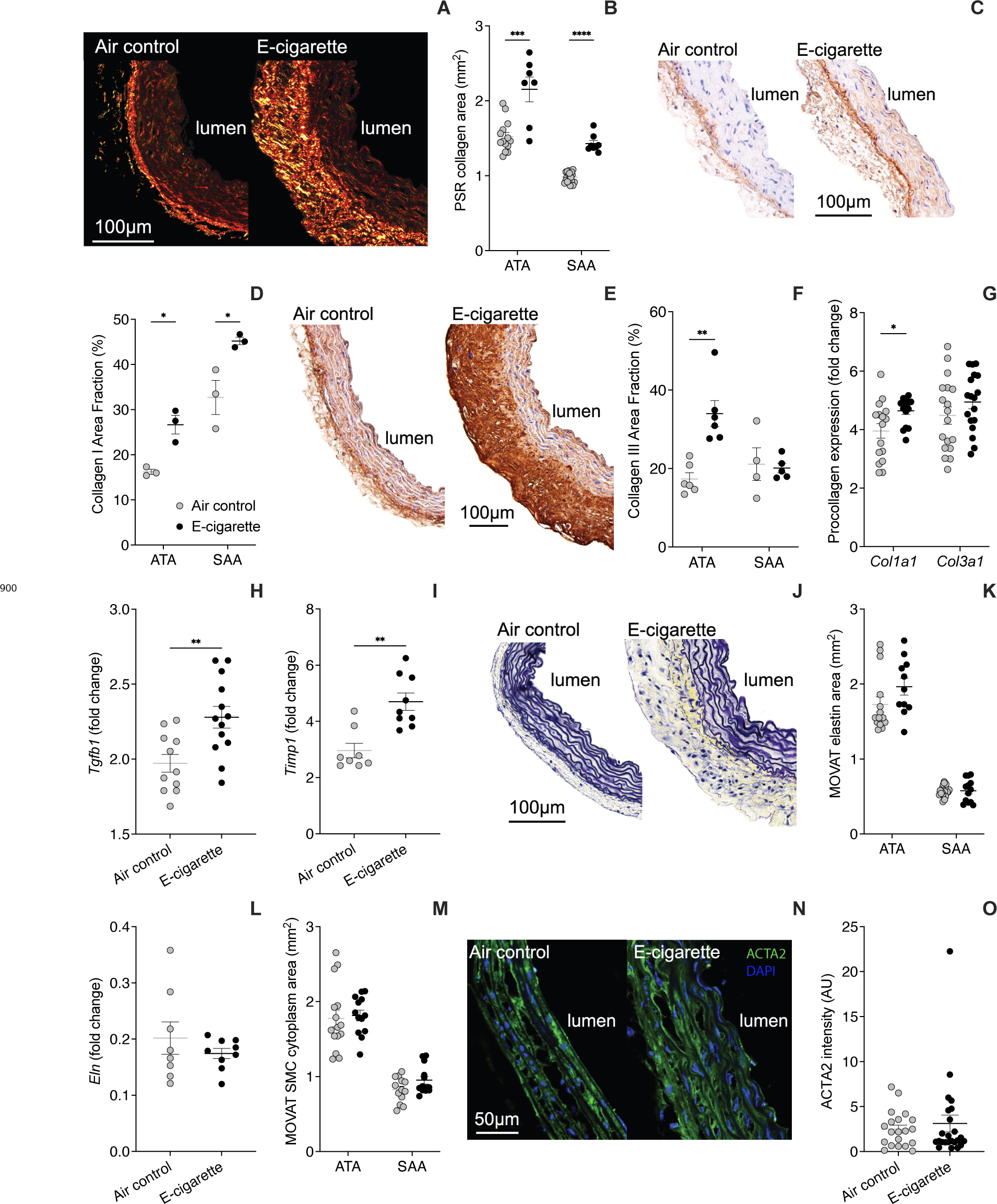
Microstructural composition of aortic tissues from air control and e-cigarette mice. Total collagen content by area (B) in aortic cross-sections stained with picrosirius red (A). Area fractions of collagen I (D) and collagen III (F) from chromogenic immunostaining (brown) of aortic cross-sections with anti-COL1A1 (C) and anti-COL3A1 (E) antibodies. mRNA expression of pro-fibrotic *Col1a1* and *Col3a1* (G), *Tgfb1* (H), and *Timp1* (I) genes in whole aorta tissue samples. Elastin (black, K) and SMC cytoplasm (purple, M) content by area in aortic cross-sections stained with MOVAT’s pentachrome (J). Expression of the *Eln* gene (L) in whole aorta tissue samples. Tissue expression of smooth muscle *α*-2 actin (ACTA2, O) from immunofluorescent staining (green) of aortic cross-sections (N). Microstructural outcomes are reported for ascending thoracic (ATA) and suprarenal abdominal (SAA) aortic tissues. Representative stains are from ATA tissues. Collagen deposition and remodeling supports thickening of the wall in the aorta of mice exposed to pod-mod e-cigarette aerosol. Statistical significance (within the same region for histology and immunostaining) denoted by * for p *<* 0.05, ** for p *<* 0.01, *** for p *<* 0.001, and **** for p *<* 0.0001 in e-cigarette *vs*. air control.

Exposure to e-cigarette aerosol did not alter the area covered by elastin in MOVAT-stained cross-sections (Figure 5J) from either ATA (e-cigarette: 1.93 *±* 0.11 mm^2^, air control: 1.72 *±* 0.10 mm^2^) or SAA (e-cigarette: 0.58 *±* 0.04 mm^2^, air control: 0.59 *±* 0.04 mm^2^) tissues (Figure 5K, Table S4), nor did it visibly affect elastic fiber integrity. As expected, *Eln* gene expression from whole aorta tissues was comparable between the ecigarette and air control groups (Figure 5L). Likewise, no difference emerged in the area occupied by SMC cytoplasm in MOVAT-stained (Figure 5J) ATA (e-cigarette: 1.82 *±* 0.07 mm^2^, air control: 1.78 *±* 0.12 mm^2^) and SAA (e-cigarette: 0.95 *±* 0.05 mm^2^, air control: 0.83 *±* 0.05 mm^2^) cross sections (Figure 5M, Table S4). Consistently, expression of smooth muscle *α*-2 actin (ACTA2, Figure 5N) was similar in aortic tissues from e-cigarette and air control mice (Figure 5O).

### Evidence of interleukin 6 (IL-6) signaling in the aorta of mice exposed to e-cigarette aerosol

Downregulation of eNOS expression, NADPH oxidase-mediated superoxide anion production, and fibrotic remodeling of the aortic wall cued surveying for evidence of pro-inflammatory IL-6 signaling and associated immune activity in response to chronic e-cigarette vaping.^41^ Plasma concentration of IL-6 in mice exposed to pod-mod e-cigarette aerosol exceeded that of air control following 8 and 16 weeks of exposure, and it remained elevated at endpoint, though not significantly so (Figure 6A). Nevertheless, the area fraction of aortic cross-sections that stained positive for vascular cellular adhesion molecule-1 (VCAM-1; Figure 6B) was larger at endpoint in e-cigarette than air control mice (Figure 6C), priming the endothelium for interactions with circulating leukocytes.

**Figure 6.**
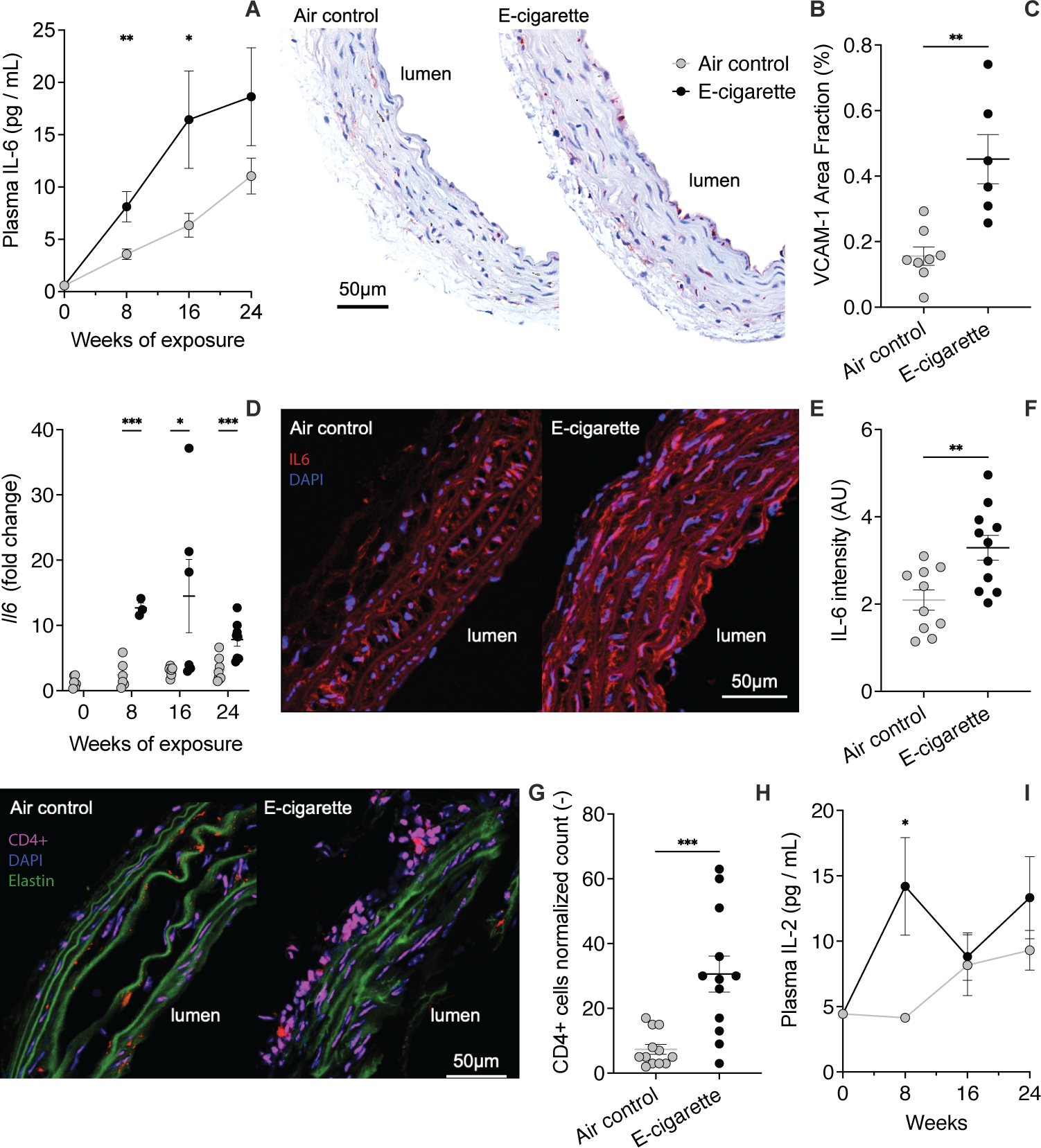
Evidence of inflammation and immune response in blood and aortic tissues from air control and e-cigarette mice. Concentration of the pro-inflammatory cytokine IL-6 in the mouse blood plasma at baseline and following 8, 16, and 24 weeks of daily exposure to either e-cigarette aerosol or filtered air (A). Expression of VCAM-1 as percent of the cross-sectional area (C) from chromogenic immunostaining (brown) of aortic tissues (B). mRNA expression of the *Il6* gene in whole aorta tissue samples at baseline and following 8, 16, and 24 weeks of daily exposure to either e-cigarette aerosol or filtered air (D). Tissue expression of IL-6 (F) from immunofluorescent staining (red) of aortic cross-sections (E). Accumulation of CD4+ T lymphocytes (H) from immunofluorescent staining of aortic cross-sections with anti-CD4 antibody (G). Concentration of the Th1-derived cytokine IL-2 in the mouse blood plasma at baseline and following 8, 16, and 24 weeks of daily exposure to either e-cigarette aerosol or filtered air (I). Pro-oxidative and pro-inflammatory signaling engages adaptive immunity in mice exposed to pod-mod e-cigarette aerosol. Statistical significance denoted by * for p *<* 0.05, ** for p *<* 0.01, and *** for p *<* 0.001 in e-cigarette *vs*. air control.

Furthermore, the expression of the *Il6* gene in the aorta of e-cigarette mice remained above air control levels following 8, 16, and 24 weeks of exposure (Figure 6D). Consistently, the mean fluorescence intensity of the IL-6 protein (Figure 6E) was higher in the e-cigarette aorta compared to air control at endpoint (Figure 6F). Finally, chronic exposure to e-cigarette aerosol prompted recruitment of CD4+ T cells (Figure 6H) throughout the aortic wall and particularly in the adventitial layer (Figure 6G). Elevation of circulating interleukin-2 (IL-2) levels beyond air control values provides additional evidence of bridging from innate to adaptive systemic immune responses following 8 weeks of e-cigarette aerosol exposure (Figure 6I).

### E-cigarette aerosol exposure did not worsen atherosclerotic lesions at the aortic root

As Apoe*^−/−^* mice spontaneously develop atherosclerosis, we sought to determine whether chronic e-cigarette aerosol inhalation perturbed the progression and severity of lesions at the aortic root. The total lesion area was comparable in MOVAT-stained aortic cross sections from pod-mod e-cigarette and air control mice (Figure 7A-B). The necrotic core likewise occupied a similar area fraction of the atheroma in e-cigarette and air control tissues (e-cigarette: 33 *±* 3%, air control: 34 *±* 3%; Figure 7C). Furthermore, neither the area occupied by collagen in atherosclerotic lesions stained with picrosirius red (Figure 7D-E) nor the thickness of the fibrous cap overlying the necrotic core (e-cigarette: 16 *±* 1 *µ*m, air control: 15 *±* 1 *µ*m, Figure 7F) were different between the e-cigarette and air control groups. Finally, exposure to e-cigarette aerosol did not affect the infiltration of CD68+ macrophages within the atheroma with respect to the air control condition (e-cigarette: 22 *±* 4%, air control: 23 *±* 6%; Figure 7G-H).

**Figure 7.**
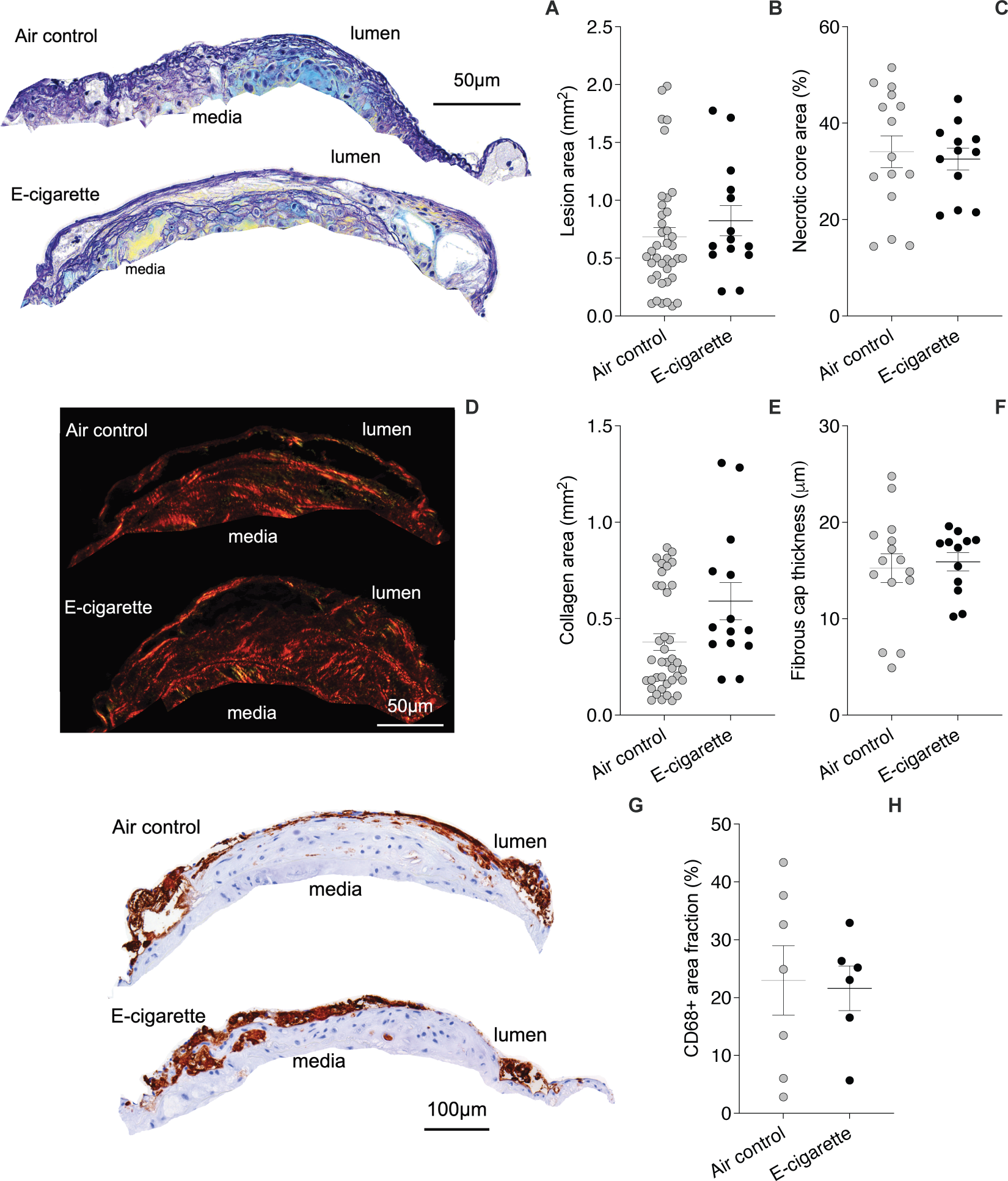
Morphology and microstructural composition of atherosclerotic lesions at the aortic root in air control and e-cigarette mice. Total lesion area (B) and percent of the total lesion area occupied by the necrotic core (C) in aortic cross-sections stained with MOVAT’s pentachrome (A). Total collagen content of the lesion by area (E) and thickness of the fibrous cap surrounding the necrotic core (F) isolated from aortic cross-sections stained with picrosirius red (D). Accumulation of macrophages as a percent of the total lesion area (H) from chromogenic immunostaining (brown) of aortic cross-sections with anti-CD68 antibody (G). Chronic exposure to pod-mod e-cigarette aerosol does not worsen lesion burden nor stability in atherosclerosis-prone Apoe*^−/−^* mice.

## 4. DISCUSSION

Pod-mod devices that aerosolize e-liquid containing nicotine salts were first introduced to the US market in 2015 and soon attracted youth e-cigarette users due to their streamlined design, ease of operation, and inconspicuous appearance.^42^ JUUL Labs, Inc.© was dominating the e-cigarette market at the time of this study and remains nowadays among the five top-selling brands.^43^ Similar to cigarette smoke^31, 44^ and prior generation (tank-style) e-cigarette aerosol,^24, 25, 28, 45^ particulates and gases produced by pod-mod devices access the body primarily through the respiratory system yet retain the ability to modulate biological processes in other organs.^46, 47^ Related to the cardiovascular system, acute inhalation of pod-mod e-cigarette aerosol increases the augmentation index (arterial stiffness), reduces the reactive hyperemia index (endothelial vasodilator function), and raises peripheral blood pressure in occasional users.^40^ Similarly, exposure to pod-mod e-cigarette aerosol causes immediate impairment of the flow-mediated dilatation (FMD) of the femoral artery in rats,^26^ perhaps due to a sympathetic autonomic reflex in response to vagal sensory nerve irritation.^16^ Encouraged by the ability of animal models to mimic the acute cardiovascular responses to pod-mod e-cigarette aerosol inhalation, we conducted a long-term exposure study in mice to evaluate whether repeated insults impart a permanent shift in baseline cardiovascular parameters, including peripheral blood pressure, endothelial function, and aortic stiffness. Our findings suggest that prolonged use of pod-mod e-cigarette products increases the cardiovascular risk by supporting oxidative stress, fibrosis, aortic stiffening, and impairment of eNOS-mediated vasodilatation, culminating in a moderate yet significant increase in systolic and pulse blood pressure.

Vasodilatation plays a crucial role in adaptive responses to physiological stimuli that require regulation of mean blood pressure and blood flow distribution. These functions are vital for maintaining vascular homeostasis and dysregulation of vasodilatory mechanisms increases the risk for cardiovascular disease. Vasodilatation occurs in response to smooth muscle cell (SMC) relaxation and depends on endothelial signaling. Although several molecules released by the endothelium are capable of inducing vasodilatation, the aorta relies primarily on nitric oxide (NO) produced by endothelial nitric oxide synthase (eNOS). Preserved vasodilatation to sodium nitroprusside (Figure 2A) indicates that aortic SMCs retain their responsiveness to NO following chronic exposure to pod-mod e-cigarette aerosol. In contrast, abated vasodilatation to acetylcholine and loss of attenuation in the acetylcholine response with L-NAME pre-incubation (Figure 2B) serve as evidence of reduced NO availability in the e-cigarette aorta.

Downregulation of both *Nos3* (Figure 2F) and eNOS protein (Figure 2D,E) expression within aortic tissues from mice exposed to pod-mod e-cigarette aerosol supports functional outcomes and suggests that blunted production may contribute to reducing NO levels. Complementary to our findings, Majid *et al.* recently reported that venous endothelial cells isolated from pod-mod e-cigarette users exhibit decreased eNOS phosphorylation in response to acetylcholine compared to never users, and that human aortic endothelial cells exposed to pod-mod e-liquids produce less NO following calcium ionophore stimulation, further implicating limited eNOS activation as a source of vaping-induced NO deficiency.^22^

Alongside reduced eNOS expression and/or activation, excess superoxide anion within the aortic wall may lower NO levels by reacting with the compound to form peroxynitrite. Paralleling observations in prior generation e-cigarette users^19^ as well as exposed mice,^28, 27^ repeated inhalation of pod-mod e-cigarette aerosol increases the oxidative burden both systemically (Figure 3G) and within aortic tissues (Figure 3A-B,F). Peroxynitrite oxidation of NO synthesis cofactor tetrahydrobiopterin (BH_4_) may in turn compel eNOS to generate additional superoxide anion instead of NO.^48^ Weakening of DHE fluorescence upon L-NAME treatment (Figure 3C,F) confirms the involvement of uncoupled eNOS in reactive oxygen species (ROS) production within aortic tissues of mice exposed to pod-mod e-cigarette aerosol, as recently reported for tank-style devices with e-liquids containing free-base nicotine.^28^

Membrane-bound NADPH oxidases (NOXs) are another critical source of ROS in the context of vaping-induced oxidative stress. Carnevale *et al.* first associated the immediate impairment of FMD in the brachial artery following e-cigarette aerosol inhalation with the decline in serum level of soluble NOX2-derived peptide, as a measure of NADPH oxidase activation in humans.^49^ Within vascular tissues, NOX2 is expressed predominantly by endothelial cells^50^ and inhibition of NOX2 reduces superoxide anion staining in the aorta of mice exposed to pod-mod e-cigarette aerosol (Figure 3E,F). Complementing our findings, both short-^27^ and long-^28^ term inhalation of e-cigarette aerosol from prior generation devices promoted expression and activation of NOX2 in the mouse aorta, while NOX2 inhibition/deficiency reduced vascular ROS production by acrolein, one of the main toxic aldehydes in e-cigarette aerosol.^27^ In contrast, NOX1 is expressed by SMCs and suppression of NOX1 activity does not relieve the oxidative burden in aortic tissues form ecigarette mice (Figure 3D,F), although NOX1 mediated superoxide anion production in rat SMCs that were directly exposed to cigarette smoke extract.^51^

Besides limiting NO availability and impeding endothelium-dependent vasodilatation, excess ROS production by NADPH oxidase has been shown to promote the fibrotic remodeling of the mouse aorta via T-lymphocyte activation, polarization, and proliferation.^52^ In line with this evidence, aortic tissues from mice exposed to pod-mod e-cigarette aerosol exhibit greater collagen content by area (Figure 5A-B, Table S4) alongside increased superoxide anion levels (Figure 3A-B,F) compared to air control. Transcriptional upregulation of the *Tgfb1* gene (Figure 5H) further suggests that TGF-*β*1 may contribute to mediating collagen deposition and preservation in response to prolonged e-cigarette aerosol inhalation. Canonic TGF-*β* signaling stimulates the synthesis of several proteins that are involved in the upkeep of the extracellular matrix, including type I collagen and tissue inhibitor of matrix metalloproteinases (TIMPs).^53, 54^ Consistently, the expression of mRNA transcripts for both the *Col1a1* (Figure 5G) and the *Timp1* (Figure 5I) genes is higher in the aorta of e-cigarette than air control mice. Offset by preserved elastin and SMC levels (Figure 5J-M, Table S4), collagen remodeling upholds wall thickening (Figure 4N, Table S3) by augmenting the cross-sectional area of the adventitia alone in the e-cigarette aorta (Table S4). Deposition of newly-synthesized collagen at increasingly larger diameters (Figure 4O, Table S3) that accommodate the progressive rise in systolic pressure (Figure 1A) may contribute to delay fiber engagement toward higher circumferential deformation in aortic tissues from e-cigarette mice (Figure 4A-B). This rightward extension of the mechanical response preserves circumferential stress (Figure 4E, Table S3) at the expense of intrinsic tissue stiffening with chronic e-cigarette aerosol inhalation, though significantly so in the abdominal segment alone (Figure 4F, Table S3). Wall thickening (Figure 4N, Table S3) and circumferential tissue stiffening (Figure 4F, Table S3) limit the relative widening of the aortic lumen (Figure 4P) that would otherwise occur between diastole and systole due to the rise of pulse pressure (Figure 1B) in mice exposed to pod-mod e-cigarette aerosol. The associated loss of distensibility (Figure 4M, Table S3) further emphasizes the structural stiffening of the e-cigarette aorta. Aligned with our *in vitro* inferences, a sharp increase in the surrogate stiffness metric of *in vivo* pulse wave velocity (PWV) was recorded in both female Apoe*^−/−^* and C57BL/6 mice that were exposed to aerosol from e-liquids with free-base nicotine over six^55^ and eight^24^ months, respectively. Distensibility quantifies the capacity of the aorta to expand and recoil throughout the cardiac cycle. Distension of the aortic wall following the release of blood into systemic circulation accommodates part of the ejected volume to contain systolic pressure, while recovery of the radial expansion augments blood flow to maintain diastolic pressure and peripheral perfusion. Reduced distensibility thus accelerates the transmission of the forward pressure wave and anticipates the return of the reflected wave, leading to an increase in systolic pressure.^56^ In line with the critical role of aortic stiffness in cardiovascular function, the structural stiffening of the aorta is as an independent predictor of the risk for cardiovascular disease.^57, 58, 59, 60, 61^ Noteworthy for contextualizing the cardiovascular effects of prolonged pod-mod e-cigarette aerosol exposure, progressive dysfunction of the vascular endothelium and stiffening of the aorta that precedes the development of overt hypertension mediate the increase in cardiovascular disease risk with aging.^62, 63^ Although we did not longitudinally track the structural stiffness of the aorta throughout the study, systolic and pulse pressure gradually increased during the initial 16 weeks of exposure, before stabilizing at time-averaged steady state values 17 mmHg and 11 mmHg higher than control, respectively (Figure 1).

Oxidative stress, endothelial dysfunction, and fibrosis linked to both reduced eNOS expression or activation and enhanced superoxide anion production by NADPH oxidase are recognized downstream effects of interleukin-6 (IL-6) signaling in the vasculature.^41^ Motivated by this observation, we therefore sought evidence of IL-6 activity in blood and aortic tissues of e-cigarette mice. Daily inhalation of pod-mod ecigarette aerosol increases plasma levels of IL-6 (Figure 6A) and elevated expression of endothelial adhesion molecules in the aorta (Figure 6B,C) supports an interaction between circulating IL-6 and the endothelium. Transcriptional upregulation of *Il6* at intermediate and final exposure time points (Figure 6D) and increased IL-6 protein content in aortic tissues following 24 weeks of daily e-cigarette aerosol inhalation (Figure 6E,F) further confirm propagation of the pro-oxidative and pro-inflammatory signaling throughout the aortic wall. Evidence of increasing IL-6 secretion by endothelial and aortic smooth muscle cells upon extension to progressively larger stretches *in-vitro*^64, 65^ suggests that the rise in cyclic circumferential deformation (Figure 4H) following e-cigarette aerosol exposure may promote IL-6 production within aortic tissues (Figure 6E,F).

Accumulation of CD4+ T-lymphocytes within the aortic wall of mice exposed to e-cigarette aerosol (Figure 6G,H) is likewise consistent with the established role of IL-6 in leukocyte recruitment, activation, and survival.^66, 67^ Although in blood rather than vascular tissues, memory and total CD4+ T-lymphocyte populations were more abundant in cigarette smokers compared to nonsmokers^68, 69, 70^ and circulating T-lymphocyte levels remained elevated in the offspring of a mouse model of prenatal cigarette smoke exposure for up to 2.5 months after birth.^71^ Note that T-lymphocytes mediate collagen remodeling and aortic stiffening due to aging alone^72, 73^ or in combination with hypercholesterolemia.^74, 75^ Despite evidence that pollutant inhalation accelerates vascular aging,^76, 77^ no study to date has examined the contribution of adaptive immunity to the structural and functional remodeling of the vasculature in the context of environmental exposure. We speculate that recruitment of CD4+ T-lymphocytes (Figure 6G,H) by IL-6 (Figure 6D-F) may support collagen secretion (Figure 5A-G) to thicken (Figure 4N) and structurally stiffen (Figure 4M) the aorta. While additional evidence is needed to corroborate this inference, findings from our study overall suggest that oxidative stress, sterile inflammation, and immune cell infiltration within the aortic wall as a result of chronic pod-mod vaping may coordinate tissue maladaptation and loss of aortic function, which increase the afterload on the heart and are thus detrimental for long-term cardiovascular health.

Notwithstanding oxidative stress and inflammation,^78, 79^ inhalation of pod-mod e-cigarette aerosol for 24 weeks did not worsen the atherosclerotic burden at the aortic root of female Apoe*^−/−^* mice compared to air controls (Figure 7). Aligned with our findings, Szostak *et al.* observed similar histopathological features in aortic root tissues from female Apoe*^−/−^* mice exposed to either e-cigarette aerosol or air for 6 months.^55^ However, both Espinoza-Derout *et al.* and Li *et al.* detected formation of larger atherosclerotic lesions at the aortic root of Apoe*^−/−^* mice upon inhalation of previous generation e-cigarette aerosol for twelve^80^ or sixteen weeks^81^ and compared to control, respectively. Exacerbation of the atherosclerotic burden at these shorter exposure time points suggests that the magnitude of the effect imposed by e-cigarette vaping on lesion composition and size may only retain significance at the early and intermediate stages of atherosclerotic disease, since lesion severity naturally progresses with age in Apoe*^−/−^* mice^37^ and most studies begin treatment around 8 weeks-of-age. Interestingly, targeted overexpression of NOX2 in the endothelium of Apoe*^−/−^* mice to enhance vascular superoxide anion production as shown here (Figure 3E-F) increased atherosclerotic disease initiation at 9 weeks-of-age, yet failed to alter lesion area at the root at later time points.^50^ In contrast, we have recently shown that cotinine-matched^12^ exposure to cigarette smoke for 24 weeks increased lesions size and promoted a vulnerable atherosclerotic phenotype in female Apoe*^−/−^* mice.^31^ Therefore, while the molecular mechanisms supporting the dynamic interaction between e-cigarette aerosol inhalation and atherosclerosis remain to be established, long-term vaping alone poses a lesser risk toward worsening advanced atherosclerotic lesions than cigarette smoking in the hypercholesterolemic milieu. Nevertheless, stiffening and inflammation of aortic tissues following chronic exposure to e-cigarette aerosol may still prime for earlier or more severe aortic disease in the presence of additional risk factors, including a high cholesterol diet and male sex that increase the propensity for atherosclerotic lesion development in Apoe*^−/−^* mice.

## 5. CONCLUSIONS

The recent introduction and rapid evolution of e-cigarette devices motivates preclinical studies in animal models of chronic exposure to infer the long-term health outcomes of vaping. We show here that daily inhalation of pod-mod e-cigarette aerosol exerts detrimental effects on the cardiovascular function of female Apoe*^−/−^* mice over the course of 24 weeks. Vaping-induced ROS production hinders endothelium-dependent vasodilatation by reducing NO availability, while tissue thickening and stiffening due to fibrotic remodeling limit the distension and recoil of the aorta between diastole and systole. Concurrent stiffening and endothelial dysfunction increase systolic blood pressure in mice exposed to e-cigarette aerosol by interfering with the capacity for adjusting aortic diameter, both passively under the effect of hemodynamic loads and actively in response to changes in blood flow. Systemic and local inflammation coupled to the engagement of adaptive immunity likely support these adverse functional outcomes that are reminiscent of vascular aging despite preserved atherosclerotic burden at the aortic root. Overall, our findings reinforce the need for imposing regulations and developing clinical guidelines regarding the use of pod-mod e-cigarette devices, to mitigate the risk for adverse cardiovascular events amongst those who vape.

## ACKNOWLEDGMENTS

The authors gratefully acknowledge the contribution of the histology cores at Tufts Medical Center and Beth Israel Deaconess Medical Center for assistance with histology and immunohistochemistry.

## SOURCES OF FUNDING

We acknowledge financial support from the National Institute of Health (R03HL142472 and R01HL168719 to CB and JMO; R01HL146627 and R01HL149927 to BR). YMF is funded by NSF GRFP fellowship (1451070). CR is funded by American Heart Association pre-doctoral fellowship (23PRE1023003).

## CONTRIBUTIONS

YMF: Methodology, Investigation, Formal Analysis, Data Curation, Visualization, Writing - Original Draft. JM: Methodology, Investigation, Writing - Review & Editing. HW, HK, CR, JV: Investigation, Writing - Review & Editing. BR: Resources, Writing - Review & Editing. JMO: Conceptualization, Supervision, Funding Acquisition, Writing - Review & Editing. CB: Conceptualization, Methodology, Supervision, Funding Acquisition, Writing - Review & Editing.

## DISCLOSURES

The authors have no disclosures.

## Notes

### Competing Interest Statement

The authors have declared no competing interest.

